# Blood-derived dietary protein promotes sleep in the mosquito *Aedes aegypti*

**DOI:** 10.1101/2025.09.24.678251

**Authors:** Jiwei Zhang, Hitoshi Tsujimoto, Samaneh Biglari, Zach N. Adelman, Alex C. Keene

## Abstract

Sleep is a ubiquitous, yet highly variable, behavior across species. The duration and timing of sleep are influenced by ecological demands and dietary context. In the mosquito *Aedes aegypti*, a blood-feeding insect with specialized nutritional requirements, the relationship between feeding and sleep remains poorly understood. Here, we investigated how blood-derived dietary protein influences sleep regulation. Using postural analysis, videography, and arousal-threshold assays, we established that immobility bouts of ≥10 minutes reliably define sleep in *Ae. aegypti*. Mosquitoes lacking the circadian clock gene *cycle* still maintained daily sleep rhythms but exhibited reduced sleep duration and heightened overall activity. Infrared activity monitoring revealed that blood-fed females showed a marked increase in sleep beginning immediately after feeding and persisting for several days, accompanied by reduced locomotor activity. Notably, this sleep elevation lasted well beyond the cessation of previously reported host-seeking phases, raising the possibility of distinct phases of opportunistic versus targeted host pursuit. To determine the dietary basis of this effect, we tested mosquitoes fed a bovine serum albumin (BSA)–based diet. BSA feeding alone was sufficient to mimic the sleep-promoting and activity-reducing effects of blood, suggesting dietary protein is a major nutritional regulator. Moreover, RNAi-mediated knockdown of the leucokinin receptor (*Lkr*), which has previously been associated with fluid homeostasis and feeding behavior, resulted in enhanced sleep and reduced activity, implicating mosquito LK signaling in the modulation of postprandial sleep. Together, these findings demonstrate that blood-derived proteins drive sustained increases in sleep and reductions in locomotor activity in *Ae. aegypti*. This work positions *Ae. aegypti* as a model for dissecting nutrient-specific regulation of sleep and highlights potential adaptive functions of protein-induced quiescence, such as energy conservation and predator avoidance. More broadly, it provides insight into how specialized diets shape the neural and behavioral architecture of sleep.

## Introduction

Sleep duration and timing varies significantly across the animal kingdom [1]. In diverse species, sleep is modified by an animal’s internal and external environment, including social cues, early-life development, stress, and food availability [2,3]. Moreover, evolutionary pressures have shaped species-specific sleep strategies, balancing the restorative benefits of sleep with ecological demands such as predation risk, foraging requirements, and reproductive behaviors [4]. While the mechanistic basis for interspecies variation in sleep remains poorly understood, there is mounting evidence that context-dependent regulation of sleep is adaptive [5,6]. Understanding how evolutionary history and cellular factors interact to regulate sleep not only has potential to provide insight into the adaptive significance of sleep but may also inform broader questions about the links between sleep, health, and survival across taxa.

Food availability potently regulates sleep [7]. Many species of animals have been shown to forgo sleep to forage under food-deprived conditions [3]. In addition, flies, fish, and mammals all display post-prandial sleep, demonstrating that sleep is bidirectionally modified by food consumption [8–10]. In insects, both total caloric content and specific dietary nutrients have been shown to influence sleep [11–13]. While this phenomenon is highly conserved, most studies have been performed in *Drosophila*, a dietary generalist, and far less is known about how diet contributes to sleep regulation in animals that consume more specialized diets [14].

Blood-feeding insects provide a unique opportunity to investigate the effects of diet on behavior [15]. Their obligate dependence on vertebrate blood imposes distinct nutritional constraints that differ markedly from the diets of generalist feeders. In addition, the dramatic physiological changes that occur following a blood meal, such as gut distension and shifts in metabolic state, are likely to influence neural circuits that govern sleep and activity [16]. The mosquito, *Aedes aegypti*, is a model for understanding the effects of blood feeding on behavior [17,18]. These insects rely exclusively on blood to complete their reproductive cycle, making them an ideal system to study the behavioral consequences of this specialized diet. While sleep has been described in this system, the effects of blood feeding on sleep regulation have not been investigated [19,20].

There is evidence to suggest that blood feeding and dietary protein influence sleep in *Ae. aegypti*. Following a blood meal, females suppress both host seeking and feeding for several days [21]. Moreover, an artificial protein-based diet composed of γ-globulins, hemoglobin, and albumin is sufficient to stimulate egg production [22], revealing dietary protein as a critical factor in reproduction and in the suppression of feeding. This suggests that dietary proteins, rather than other components of blood, underlie these behavioral shifts. Nevertheless, the specific amino acids that mediate the regulation of reproductive and feeding behaviors remain unidentified. Examining sleep across different feeding conditions therefore provides an opportunity to determine which dietary macronutrients regulate sleep in *Ae. aegypti* and to test whether these macronutrients overlap with those that regulate other blood-feeding–associated behaviors.

Here, we examine sleep behavior in *Ae. aegypti* and tested the effects of blood feeding and dietary protein on sleep. We found that both blood meals and a bovine serum albumin (BSA)-based protein diet increase sleep for several days. Critically, we found that the period of increased sleep is substantially longer than the cessation of host seeking, suggesting the existence of discreet opportunistic vs active host-seeking phases. These findings establish *Ae. aegypti* as a model for investigating the effects of diet on sleep regulation and imply that methods to disrupt mosquito host-seeking must consider both of these behavioral systems.

## Results

Sleep has previously been defined in three mosquito species as inactivity bouts that are greater than two hours based on postural analysis [20]. We sought to further define periods of immobility used to define sleep using multiple established criteria including arousal threshold, posture, and circadian regulation [23]. To identify species-specific postures during the day and night, individual mated female mosquitos were recorded at high resolution. Key postural differences were identified with inactive and active mosquitoes that were consistent with those previously used to define sleep [20]. Shortly following activity, the hind legs extend upward while the proboscis points downward, often contacting the ground revealing a posture associated with waking activity (Figure 1A–B). Following immobility lasting on average 20 min during the day, and 11 min in the evening, the hind legs are in fully contact with the tube and the proboscis becomes elevated (Figure 1C–E). These findings suggest a shorter duration of immobility is associated with sleep than has previously been reported.

**Figure 1.**
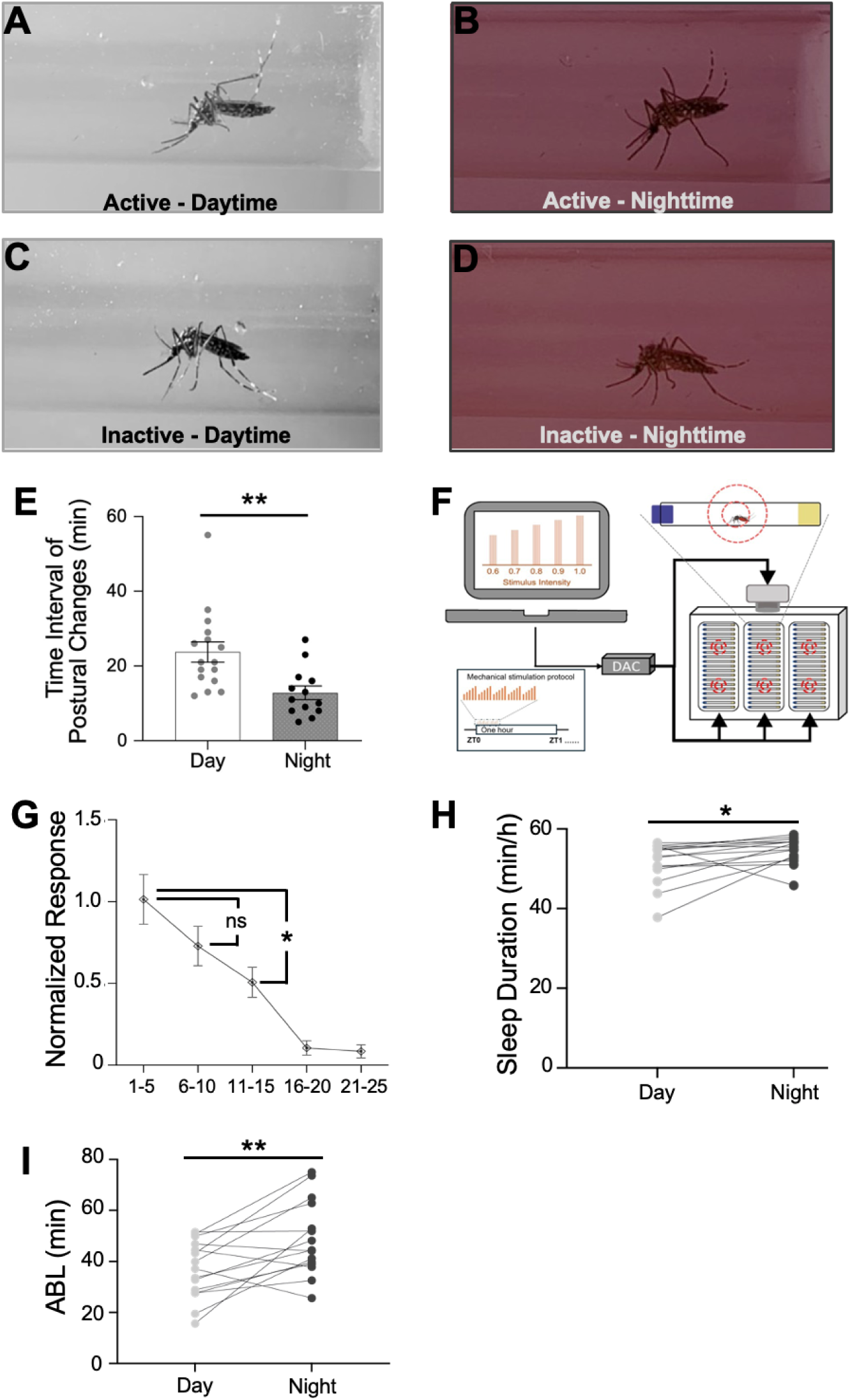
Characterization of sleep and arousal threshold in female *Ae. aegypti*. (**A-D**) Representative images of female *Ae. aegypti* either in active state or during sleeping. These images illustrate postural differences of the hind legs in active or inactive states, which were recorded via videography under the daytime or nighttime. (**E**) Quantification of averaged time interval that individual animal required to change postures. Averaged time interval to shift postures was significantly longer during the day (clear bar) than that at night (gray-filled bar). Data were assessed for normality using Shapiro-Wilk test. As the Day group violated the normality assumption (*P* < 0.05), comparisons between Day and Night groups were analyzed using a two-tailed Mann-Whitney U test: *P* = 0.0021 and U = 174.50. Individual data points are shown as gray or black for selective bouts during daytime (N=16) or black for bouts during nighttime (N=13). **(F)** Schematic of modified *Drosophila* ARousal Threshold (DART) system using for assessment of sleep responsiveness in mosquitoes. The mechanical stimuli were delivered once per hour for one LD cycle, beginning at ZT0, with protocol shown in bottom left box. **(G)** Behavioral threshold assessment in mosquitoes across a single light-dark cycle (LD, 15 h:9 h) using a modified DART system. To determine the behavioral threshold, mosquitoes were subjected to vibratory stimuli of incrementally increasing intensity [0.6g–1.0g (acceleration)] once per hour throughout one complete LD cycle. For each stimulation event, the duration of the ongoing inactive bout prior to stimulation was recorded. Responses were normalized to the mean response within each stimulus intensity bin, such that values represent proportional changes relative to the individual baseline. The data indicated that mosquitoes exhibiting shorter periods of inactivity (1–9 min) frequently responded to the stimulus, whereas those in longer inactive bouts (≥10 min) were significantly less responsive, which suggested the onset of deeper sleep following approximately 10 minutes of continuous inactivity, thereby validating the use of a 10-minute immobility threshold for defining sleep behaviorally in this context. One-way ANOVA F_4,140_ = 14.93, (1-5 min) vs. (6-10 min) *P* = 0.2460; (1-5 min) vs. (11-15 min) *P* = 0.0451. N=29. (**H-I**) Individual mosquito trajectories showing increased sleep duration (**H**, paired t test, *P* = 0.0478) and averaged bout length (ABL) during nighttime (**I**, paired t test, *P =* 0.0041). Error bars represent the standard error of the mean (±SEM). Here and in subsequent figures, asterisks indicate the level of significance: \**P* < 0.05, \*\**P* < 0.01, \*\*\**P* < 0.001.

To define further the period of immobility associated with sleep in *Ae. Aegypti*, we tested mosquitoes in the *Drosophila* ARousal Threshold (DART) that has been widely used to measure sleep duration and intensity in flies (Figure 1F) [24–26]. The responsiveness to stimuli was significantly reduced at periods of 10 minutes of immobility or longer (Figure 1G). Sensory responsiveness decreased further for bout lengths greater than 15 min, raising the possibility of light and deep sleep, or that a fraction of the animals is asleep at 10 minutes. Together, these findings suggest that bout lengths of 10 min or longer can be used to define sleep. Using the 10 min of immobility to define sleep, we then sought to determine whether sleep is under circadian regulation. Periods of immobility were quantified using videography obtained in the DART system. Both total sleep duration, and the average length of each sleep bout were greater in the nighttime as compared to the daytime (Figure 1H–I). Therefore, behavioral posture, and arousal threshold, and circadian regulation support a 10 min immobility threshold in *Ae. aegypti*.

Our findings, and those of others, suggest *Ae. aegypti* are diurnal [27,28]. The circadian clock influences both sleep duration and the timing of sleep in *Drosophila* and mouse [29–31]. To determine the effects of the circadian clock in sleep in *Ae. aegypti,* we compared sleep in mosquitos lacking the core circadian gene *cycle* (*AeCyc^−/−^*) to wildtype (WT) control mosquitoes [32]. We applied infrared-based activity monitors that have been widely used to measure sleep in *Drosophila,* as well as *Ae. aegypti*, to analyze sleep [20,33]. Total activity was elevated in *AeCyc^−/−^* mosquitoes compared to wildtype during the day and night periods (Figure 2A–B). The dusk activity peak was also largely lost in *Cyc^−/−^*mosquitoes, suggesting the circadian clock is required for regulation of daily activity, even under light-dark condition (Figure 2C). While we assessed the loss of *Cyc* on sleep regulation, sleep was significantly reduced during the day and night periods in *Cyc^−/−^* mosquitoes, suggesting that *Cyc* promotes sleep (Figure 2D–E). Nevertheless, sleep in both wildtype and *Cyc^−/−^*mosquitoes was greater during the nighttime than the daytime. We applied a Markov model that determines propensity to remain asleep, which serves as an indicator of sleep depth [34]. The wake propensity was greater in *AeCyc^−/−^*while sleep propensity was diminished (Figure 2F–G). Together, these findings suggest that *Cyc* modulates the timing of locomotor activity and the duration of sleep.

**Figure 2.**
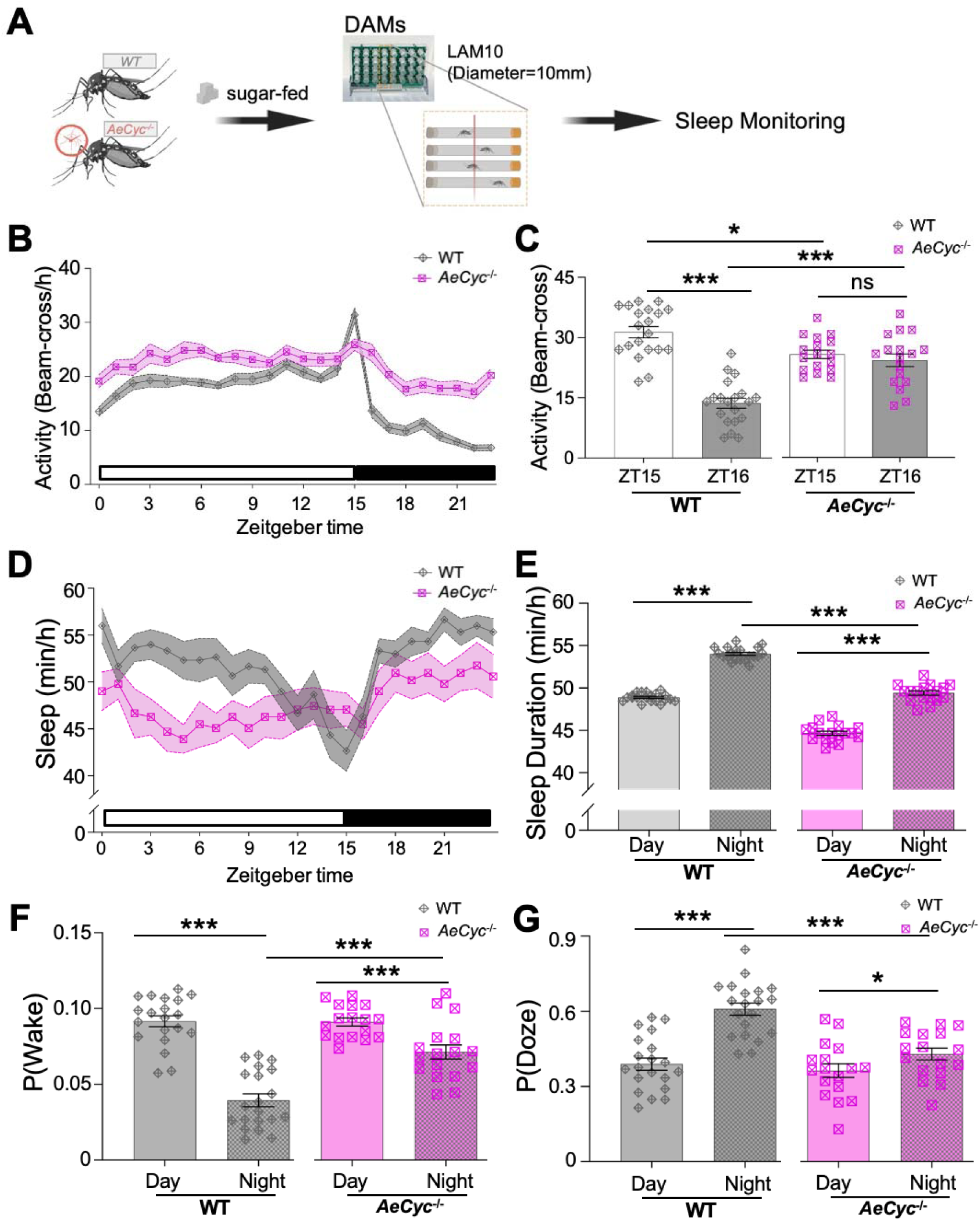
Reduced day and night sleep duration in *AeCyc^−/−^* mosquitoes. **(A)** Schematic of sleep monitoring in wide type (WT) and *AeCyc^−/−^* mosquitoes via infrared-based LAM10 (tube outside diameter = 10 mm, length = 100 mm). **(B)** Activity profile across day and night cycles (15 h:9 h) for WT (gray) and *AeCyc^−/−^*(pink) mosquitoes. *AeCyc^−/−^* mutants showed elevated activity during both day and night periods compared to WT animals. Light-colored shadows indicate the ±SEM error bar, and white and black boxes indicate the daytime and nighttime, respectively. (**C**) Two-way ANOVA revealed a significant interaction between genotype and ZT time (F_1,70_ = 36.56, *P* < 0.0001). WT mosquitoes showed a sharp decrease in activity from ZT15 to ZT16 (paired t-test, *P* < 0.0001), whereas *AeCyc^−/−^* mutants maintained high activity levels at ZT16 (paired t-test, *P* = 0.4084). At ZT15, *AeCyc^−/−^* activity was significantly lower than WT (unpaired t-test, *P = 0.0045*, Cohen’s d = 1.00), while at ZT16 *AeCyc^−/−^* activity was significantly higher than WT (unpaired t-test, *P* < 0.0001, Cohen’s d = 1.78), suggesting the dusk activity peak was largely lost in *AeCyc^−/−^* mosquitoes. **(D)** Sleep profile across day and night cycles for WT (gray) and *AeCyc^−/−^* (pink) mosquitoes. *AeCyc^−/−^*mutants showed decreased sleep during both day and night periods compared to WT animals. **(E)** Two-way ANOVA revealed significant main effects of genotype (F_1,70_ = 515.66, *P* < 0.0001) and time of day (F_1,70_ = 653.43, *P* < 0.0001), but no significant interaction (F_1,70_ = 0.98, *P* = 0.3261). Both WT and *AeCyc^−/−^*mosquitoes slept more during the night than during the day (paired t-test, WT: *P* < 0.0001; *AeCyc^−/−^* : *P* < 0.0001). However, *AeCyc^−/−^* mutants exhibited significantly reduced sleep duration compared to WT controls during both the day (Welch’s t-test, *P* < 0.0001) and the night (unpaired t-test, *P* < 0.0001). **(F)** Two-way ANOVA revealed significant main effects of genotype (F_1,70_ = 16.23, *P* = 0.0001) and time of day (F_1,70_ = 92.29, *P* < 0.0001), with a marginally significant interaction (F_1,70_ = 17.35, *P* < 0.0001). Both WT and *AeCyc^−/−^* mosquitoes showed higher P(Wake) during the day than at night (WT: Wilcoxon signed-rank test, *P* < 0.0001; *AeCyc^−/−^*: paired t-test, *P* = 0.0021). Notably, *AeCyc^−/−^* mutants exhibited a significantly elevated P(Wake) during the night compared to WT controls (Mann-Whitney U test, U = 41.00, *P* < 0.0001), whereas daytime P(Wake) did not differ between genotypes (unpaired t-test, *P* = 0.9081). These results indicated that *AeCyc^−/−^* was specifically required for maintaining low wake probability during the night. **(G)** Two-way ANOVA revealed significant main effects of genotype (F_1,70_ = 16.80, *P* < 0.0001) and time of day (F_1,70_ = 35.69, *P* < 0.0001), with a significant interaction (F_1,70_ = 9.38, *P* = 0.0031). Both WT and *AeCyc^−/−^* mosquitoes showed higher P(Doze) during the night than during the day (WT: paired t-test, *P* < 0.0001; *AeCyc^−/−^*: paired t-test, *P* = 0.0428). *AeCyc-/-* mutants also exhibited a significantly reduced P(Doze) during the night compared to WT controls (unpaired t-test, *P* < 0.0001). Error bars represent the standard error of the mean (±SEM). Data points in the bar graphs represent individual animals; N=20 for WT in gray and N=17 for *AeCyc^−/−^* in pink.

To determine if diet impacts sleep, we compared differences between sugar or blood fed females. Female mosquitoes aged 3-5 days after eclosion were blood-fed with defibrinated sheep blood, and after three days were allowed to lay eggs (Figure 3A). We focused on the period after oviposition as we were primarily interested in behavioral differences induced by blood feeding that were independent of host-seeking, which is restored by 72 h after feeding [35]. Following oviposition, mosquitoes were lightly anesthetized on ice and loaded into custom-built infrared-based activity monitors, like those regularly used to measure sleep and activity in fruit flies. A sleep profile beginning on day 4 following feeding revealed persistently elevated sleep for blood-fed mosquitoes across the day and night periods (Figure 3B–C). Analysis over time revealed that sleep was significantly elevated in blood-fed mosquitoes for Days 4 and 5, before returning to sugar-fed levels (Figure 3D) [33]. On Day 4 following blood feeding, mosquitoes displayed reduced wake propensity and increased sleep propensity (Figure 3E–F). These findings suggest blood feeding induces long-lasting increases in sleep that extend far beyond the window where host responsiveness/seeking is restored.

**Figure 3.**
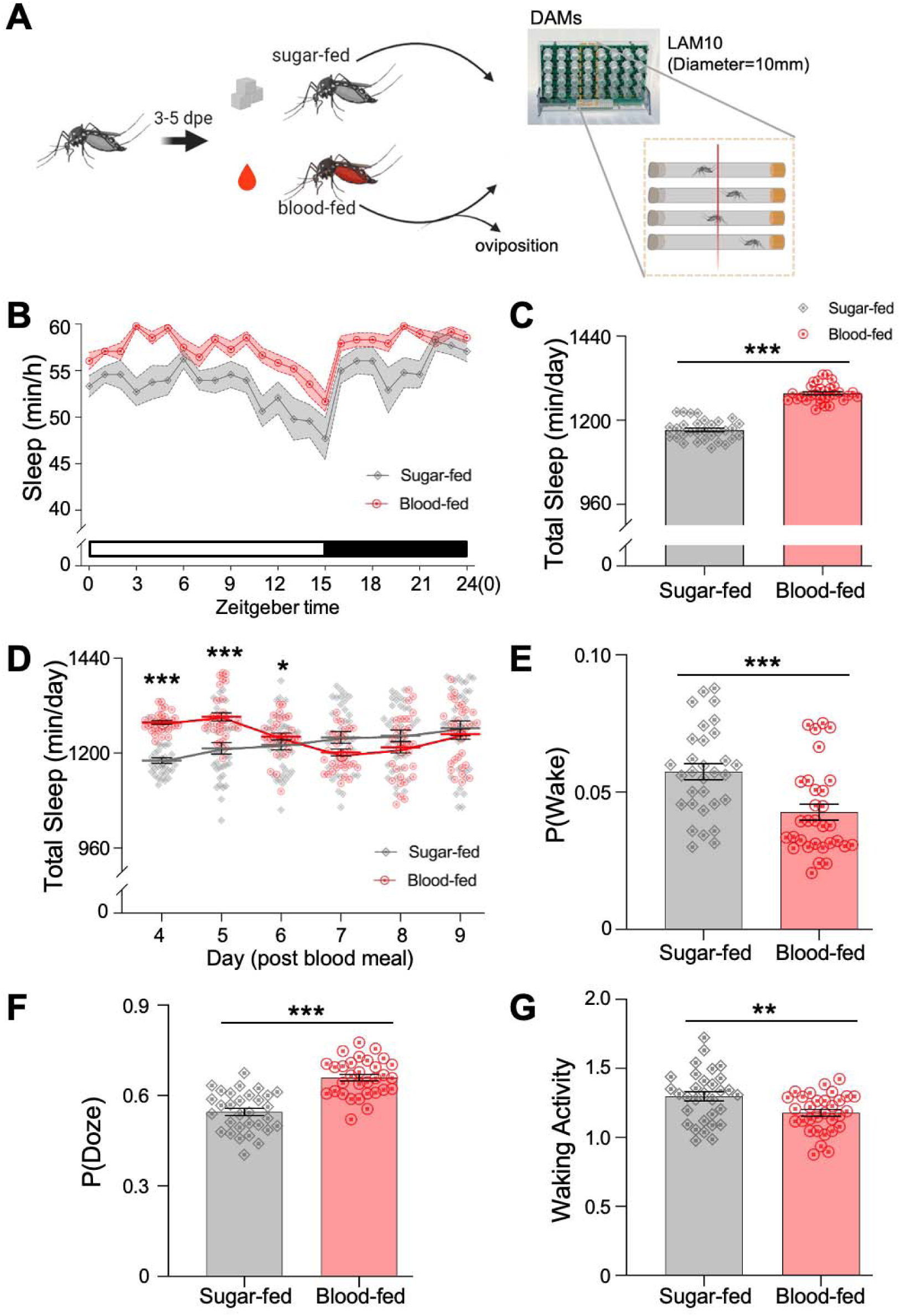
Effects of blood feeding on sleep patterns in *Ae. aegypti*. **(A)** Schematic representation of the experimental design. Female mosquitoes, aged 3-5 days post-eclosion (dpe), were either sugar-fed or blood-fed. Following oviposition, individual animals were loaded to the infrared-based activity monitors (LAM10, diameter=10 mm) used for tracking sleep and activity. (**B)** Sleep profile (min per hour) across the 24 h cycle on Day 4 post-oviposition. Blood-fed mosquitoes (red line, N=31) displayed a consistently elevated sleep profile compared to sugar-fed counterparts (gray line, N=31). Light-colored shadows indicate the ±SEM error bar, and white and black boxes indicate the daytime and nighttime, respectively. (**C**) Comparison of total sleep of blood-fed mosquitoes (red cycles) on Day 4 post blood meal to sugar-fed controls (gray diamonds). Blood-fed mosquitoes showed a significant increase in total sleep compared to sugar-fed mosquitoes (Unpaired t-test, *P* < 0.0001). (**D**) Total sleep duration across Day 4 to 9 post-blood meal. Two-way ANOVA revealed significant effects of time, feeding status, and their interaction (all *P* < 0.0001). Blood-fed mosquitoes exhibited significantly increased sleep compared to sugar-fed controls on Days 4-6 (Day 4: *P* < 0.0001; Day 5: *P* < 0.0001; Day 6: *P* = 0.021), but not on Days 7-9. (**E-G)** Changes of sleep architecture of blood-fed mosquitoes (red cycles) on Day 4 post blood meal compared to sugar-fed controls (gray diamonds). Blood-fed mosquitoes demonstrated a significantly lower waking propensity, P(Wake), compared to sugar-fed mosquitoes (**E**, Mann-Whitney U test = 726, *P =* 0.0006). Blood-fed mosquitoes exhibited a significantly higher sleeping propensity, P(Doze), compared to sugar-fed controls on Day 4 post-blood feeding (**F**, Unpaired t-test, *P* < 0.0001). Error bars represent the standard error of the mean (±SEM). Data points in the bar graphs represent individual animals. Waking activity (activity per waking minute) was reduced on Day 4 post blood meal compared to that of sugar-fed animals, indicating lower movement intensity after blood feeding (**G**, Unpaired t-test, *P* = 0.0063).

In *Drosophila,* periods of sleep for 30 minutes of longer are associated with deep sleep[36]. Using this analysis, blood fed mosquitos slept significantly more than sugar-fed mosquitoes, validating the findings using the previously identified 10-minute-immobility definition (Figure 3–figure supplement 1). Therefore, we sought to reanalyze the effects of blood meals using longer sleep bouts. To determine whether diet induces hypoactivity independent from sleep, we normalized activity to account for differences in sleep between conditions, as has previously been described in fruit flies [33,37]. The waking activity (activity per waking minute) was reduced compared to sugar fed controls, indicating less intensity of movement (Figure 3G). Analysis through nine-days post-blood feeding revealed that waking activity was reduced in blood-fed mosquitoes through Day 5, before returning to normal (Figure 3–figure supplement 1). Therefore, blood fed mosquitoes are hypoactive for a time course that mirrors the sleep increases [24].

To further define the time-course associated with dietary changes in sleep and determine whether changes occur prior to oviposition, we sought to measure sleep immediately following a blood meal. Mosquito behavior was recorded in 6-well tissue culture plates containing 1% agar and 10% sucrose for the first day following blood feeding (Figure 4A), and behavior was then tracked using EthoVision XT system as we have previously described in *Drosophila* [33]. An increase in sleep and diminished velocity profile was detected immediately after blood feeding and this persisted over the 24 h recording (Figure 4B-C; and Figure 4–figure supplement 1). This was associated with reduced total locomotor activity (Figure 4D-E). These findings confirm that blood feeding has both immediate and prolonged effects on sleep and locomotor activity. To more closely examine the effects of blood feeding on sleep and activity, we sought to extract measures of sleep intensity from video-tracking data. This analysis revealed reduced P(Wake) and increased P(Doze) in blood-fed mosquitoes, confirming that blood feeding increases sleep propensity (Figure 4–figure supplement 1).

**Figure 4.**
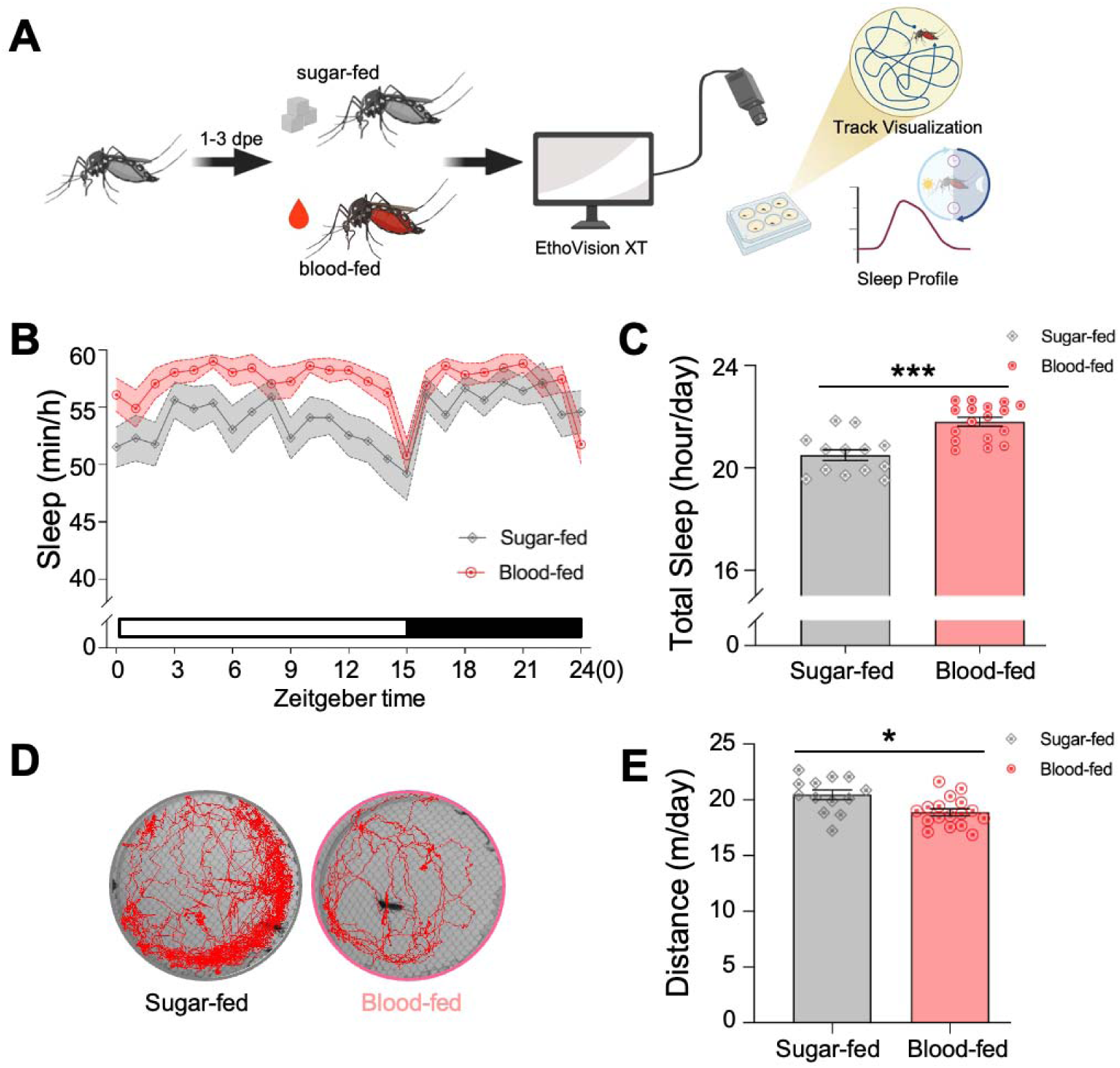
Validation of blood feeding-induced sleep in female *Aedes aegypti* using EthoVision XT. (**A**) Schematic of the EthoVision XT-based behavioral recording assay. Female mosquitoes at 1–3 days post-eclosion (dpe) were provided with either a sugar meal or a blood meal and then individually transferred into a 6-well plate immediately after feeding for continuous behavioral monitoring over one LD cycle. EthoVision XT was used for automated locomotor tracking, trajectory visualization, and sleep analysis. (**B**) Sleep profiles across the 24 h LD cycle in sugar-fed and blood-fed mosquitoes. Blood-fed females exhibited increased sleep relative to sugar-fed controls during both the light and dark phases. Lines represent mean values, and shaded areas indicate ± SEM. White and black bars denote the light and dark phases, respectively. (**C**) Total daily sleep duration in sugar-fed and blood-fed mosquitoes. Blood-fed females showed a significant increase in total sleep duration compared with sugar-fed controls (unpaired t-test, *P* < 0.0001). (**D**) Representative locomotor trajectories of sugar-fed and blood-fed mosquitoes recorded by EthoVision XT. Red lines indicate movement paths, illustrating reduced activity in blood-fed females. (**E**) Blood feeding significantly reduced total locomotor distance compared to sugar feeding (unpaired t-test, *P* = 0.015), reflecting the concomitant increase in sleep behavior. Gray denotes sugar-fed mosquitoes (N=13) and red denotes blood-fed mosquitoes (N=17). Error bars represent ± SEM, and each dot in the bar graphs represents one individual mosquito.

Protein represents a primary dietary component that contributes to sleep regulation [11,38]. Feeding mosquitoes bovine serum albumin (BSA), with ATP included as a phagostimulant, has previously been shown to induce egg laying, indicating that proteins represent a blood-derived dietary cue that modulates behavior [39]. To determine whether proteins are sufficient to increase sleep, we compared sleep in sugar-fed and BSA-fed mosquitoes beginning on Day 4 after feeding, using the same post-feeding time course applied in the blood-feeding experiments (Figure 5A). Sleep was elevated in BSA-fed mosquitoes on Day 4 following feeding compared to sugar-fed controls and was persistently elevated during the day and the night periods (Figure 5B–C). Sleep measurements over six days revealed that sleep in BSA-fed mosquitoes was elevated for Day 4 and 5, then returned to sugar-fed levels (Figure 5D). The increase in sleep duration was associated with reduced wake propensity and increased sleep propensity supporting deeper sleep (Figure 5E–F). Furthermore, waking activity was reduced in BSA-fed animals, suggesting that protein contributes to hyperlocomotion (Figure 5G). Together, these findings reveal that the BSA feeding phenocopies the sleep-promoting and hypoactive effects of blood feeding, suggesting it is specifically dietary protein that promotes sleep.

**Figure 5.**
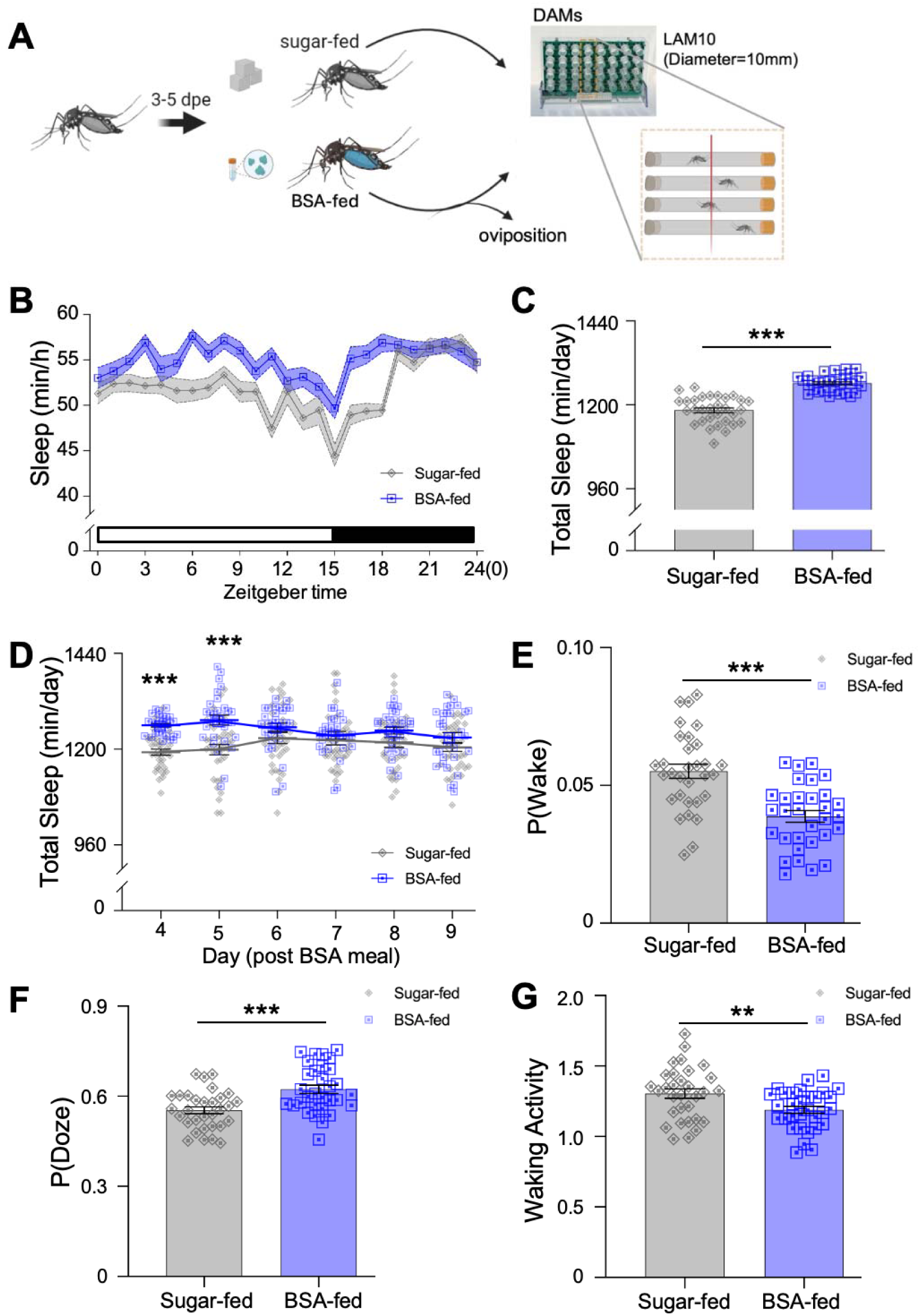
Dietary protein increases sleep in female *Ae. aegypti* via BSA feeding. **(A)** Schematic overview of the experimental design. Female mosquitoes, aged 3-5 dpe, were either sugar-fed or fed Bovine Serum Albumin (BSA). Following oviposition, sleep was monitored using custom-built infrared activity monitors over a six-day recording period. **(B)** Comparison of total sleep of BSA-fed mosquitoes (blue squares) on Day 4 post BSA meal to sugar-fed controls (gray diamonds). Total sleep duration of BSA-fed mosquitoes showed a significant increase compared to sugar-fed mosquitoes (Welch’s t-test, *P* < 0.0001). (**C**) Sleep distribution across one LD 15 h:9 h cycle on Day 4 post BSA meal. BSA-fed mosquitoes (blue line) demonstrated elevated sleep levels during both day and night periods compared to sugar-fed mosquitoes (gray line). Light-colored shadows indicate the ±SEM error bar, and white and black boxes indicate the daytime and nighttime, respectively. (**D**) Total sleep duration across Days 4 to 9 post BSA feeding. Two-way ANOVA revealed significant effects of time, feeding status, and their interaction (*P* = 0.009). BSA-fed mosquitoes (blue) exhibited significantly increased sleep compared to sugar-fed controls (gray) on Days 4 and 5 post BSA feeding (Day 4: Welch’s t-test, *P* < 0.0001; Day 5: unpaired two-tailed t-test, *P* = 0.0006), but not on Days 6-9 (Day 6: *P* = 0.252; Day 7: *P* = 0.526; Day 8: *P* = 0.130; Day 9: *P* = 0.317). **(E-G)** Changes of sleep architecture of BSA-fed mosquitoes (blue) on Day 4 compared to sugar-fed mosquitoes (gray). Significantly lower P(Wake) (**E**, unpaired t-test, *P* < 0.0001) and higher P(Doze) (**F**, unpaired t-test, *P* = 0.0003) was showed in BSA-fed population compared to sugar-fed group on Day 4 post-BSA feeding. BSA-fed mosquitoes also exhibited reduced waking activity compared to sugar-fed controls (**G**, unpaired t-test, *P* = 0.0082). Each dot in the bar graphs represents one individual mosquito, and blue squares indicate BSA-fed mosquitoes (N=31) and gray diamonds sugar-fed individuals (N=31).

The *Ae. aegypti leucokinin receptor* (*Lkr*) (VectorBase ID: AAEL006636), has previously been shown to modulate fluid excretion following a blood meal and sugar perception [40–46], and has been implicated in regulating sleep in *Drosophila* [47], raising the possibility that it also impacts sleep in mosquitoes. To test for the role of LK signaling in sleep regulation, we targeted *Lkr* in blood fed animals and measured the effects on sleep. Briefly, mosquitoes were blood fed, then injected with dsRNA targeted to either *Lkr* or *EGFP* and tested for sleep immediately following injection and oviposition period (Figure 6A). Surprisingly, loss of *Lkr* increased sleep over control mosquitoes injected with dsRNA targeted to *EGFP* for Days 4 and 5 following blood feeding (Figure 6B–D). This was accompanied by a reduction in wake propensity and an increase in sleep propensity (Figure 6E–F). Furthermore, waking activity was reduced in *Lkr* knockdown mosquitoes compared to control mosquitoes (Figure 6G). Therefore, silencing *Lkr* promotes sleep and reduces locomotion similarly to blood feeding or BSA feeding. These findings raise the possibility that blood-feeding inhibits LK signaling to promote sleep.

**Figure 6.**
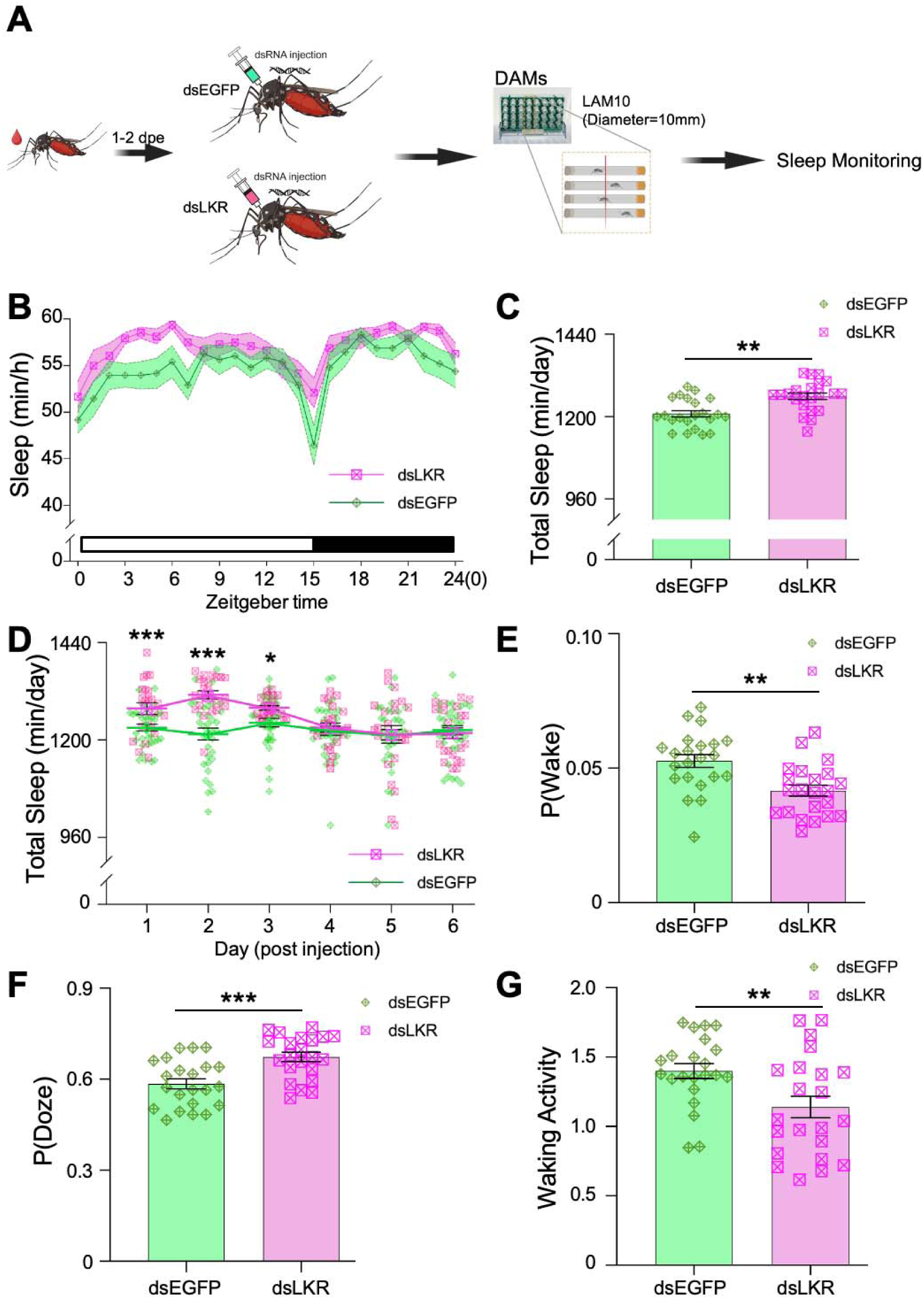
Role of the leucokinin receptor (*Lkr*) in sleep regulation of female *Ae. aegypti*. (**A**) Schematic overview of the experimental design for sleep monitoring in dsRNA-injected mosquitoes. (**B**) Sleep profiles of female mosquitoes injected with either *dsLKR* or *dsEGFP* on Day 1 post injection. Mosquitoes with *Lkr* knockdown (pink) displayed a distinct sleep pattern, with increased sleep observed during both daytime and nighttime periods compared to *dsEGFP* group (green). Light-colored shadows indicate the ±SEM error bar, and white and black boxes indicate the light and dark periods, respectively. (**C**) Daily sleep duration was significantly induced in *dsLKR*-injected mosquitoes (pink squares) compared to *dsEGFP* controls (green diamonds) (unpaired two-tailed t-test, *P* = 0.0004). (**D**) Total sleep duration across Days 1 to 6 post-siRNA injection. Mosquitoes injected with *dsLKR* (red line with pink squares) exhibited significantly increased sleep compared to controls injected with *dsEGFP* (green line with diamond markers) on Days 1-3 (Day 1: unpaired t-test, , *P* = 0.0004; Day 2: Welch’s t-test, *P* = 0.0005; Day 3: Welch’s t-test, *P* = 0.012), but not on Days 4-6 (Day 4: *P* = 0.184; Day 5: *P* = 0.511; Day 6: *P* = 0.477). (**E**-**G**) Comparisons of P(Wake), P(Doze) and waking activity of mosquitoes injected with *dsLKR* to *dsEGFP* controls on Day 1 post injection. After knockdown of *Lkr*, female mosquitoes demonstrated a significantly lower probability of waking compared to *dsEGFP* injected controls (**E**, unpaired t-test, *P* = 0.0011). In contrast, the sleeping propensity, P(Doze), was significantly induced in *dsLKR*-injected mosquitoes compared to *dsEGFP* controls (**F**, unpaired t-test, *P* = 0.0003). *dsLKR* injected mosquitoes exhibited significantly reduced waking activity compared to *dsEGFP* controls (**G**, unpaired t-test with Welch’s correction, *P* = 0.0094). Error bars represent the standard error of the mean (±SEM), and asterisks indicate the level of significance: \**P* < 0.05, \*\**P* < 0.01, \*\*\**P* < 0.001. Each dot represents one individual mosquito, and green diamonds indicate *dsEGFP*-injected mosquitoes (N=22) and pink squares with cross makers indicate individuals injected with *dsLKR* (N=22).

## Discussion

Standardized behavioral characteristics of sleep including elevated arousal threshold, rebound following deprivation and species-specific posture, have been used to characterize sleep in species ranging from the jellyfish *Cassiopeia* to humans [4,23,48]. Sleep can be characterized by physiological changes in brain activity or by monitoring the behavioral correlates that accompany these changes [48,49]. Here, we apply posture analysis, videography, and measurement of arousal threshold to define sleep as periods of immobility lasting greater than 10 minutes. This is longer than the five minutes of immobility that is widely used to define sleep in *Drosophila*, and shorter than the 120 minutes that has previously been used to define sleep in *Ae. aegypti* [20,50]. While the 10-minute definition appears to meet existing criteria, mosquitos displayed a higher arousal threshold at 15 minutes raising the possibility of multiple sleep states, or that 10 minutes of immobility only results in a subset of animals being asleep. Therefore, additional analysis may be helpful in refining the precise amount of inactivity associated with sleep.

Additional approaches have been used in other models to refine the definition of sleep. For example, indirect calorimetry has been applied in flies and rodents to measure sleep-associated changes in whole-body metabolic rate [51,52]. Application of this approach could define whether the graded changes in arousal threshold observed in mosquitoes represent light and deep sleep. In addition, analysis of microbehaviors such as haltere movements associated that are associated with sleep in flies could be applied to mosquitoes to better define sleep duration and quality [53,54]. Finally, sleep in flies is associated with changes in local-field potentials and neural activity that could be measured in mosquitoes [55,56]. Therefore, while our findings support the use of 10 minutes of immobility to define sleep, the application of additional methods will further improve our understanding of sleep quality. In addition, it would be particularly valuable to apply these analyses to *AeCyc^−/−^*mosquitoes with shortened sleep, as well as future genes identified to impact sleep, in order to determine the effects of these genetic mutations on sleep depth and sleep-associated changes in neural activity.

While we have focused on behavioral analysis of sleep, both molecular and physiological changes have been used to define sleep in *Drosophila.* For example, sleep is associated with a reduction in synaptic proteins throughout the brain, providing a molecular readout of sleep need [57]. In addition, sleep can be identified electrophysiologically through slow-wave oscillations in sleep regulatory networks, through 7-10HZ oscillations in local field potentials, or through wide-spread reductions in network connectivity imaged using Ca^2+^ sensors [55,58,59]. A number of groups have used GCaMP-expressing mosquitos to measure various aspects of brain activity including chemosensation, and host-seeking behavior [60–62]. The application of these approaches to study sleep could help further define the behavioral and physiological underpinnings of sleep and wakefulness.

Here, we identify a critical role for dietary protein in regulation of mosquito sleep. A central question is whether specific protein components within blood promote sleep. Prior work in fruit flies has shown that tryptone protein alone is sufficient to induce post prandial sleep [11]. In addition, feeding flies the amino acids D-Serine or D-Glutamine alone increases sleep, suggesting that individual amino acids are sufficient to promote sleep [63]. Studies in mosquitoes have attempted to identify dietary components associated with egg production and refeeding. An artificial protein-based diet consisting of gamma-globulins, hemoglobin and albumin was able to stimulate egg production in *Ae. aegypti,* revealing dietary protein to be a critical factor in reproduction [64–66]. One consistent theme amongst these studies is that blood plasma or BSA can trigger ecdysteriod production and vitellogenesis, but equivalent proportions of free amino acids cannot, suggesting that a diverse complement of amino acids may be required [67,68]. Applying similar approaches of minimal diets and individual amino acid supplementation may help identify the specific dietary components that promote sleep or wakefulness.

A previous study described reduced activity immediately following a blood meal, followed by increased activity post-egg laying [69]. The identified hyperactivity was associated with humidity seeking and dependent on the *cycle* gene. Conversely, our studies find prolonged reductions in activity following blood-feeding [69]. It is possible that the temporal differences in hypoactivity following feeding are due to the differences in the maintenance of mosquitos prior to, and during the assay. Therefore, further examination of the temporal dynamics associated with hypolocomotion across multiple behavioral paradigms, and particularly those that mirror the animal’s natural ecology will be important.

Genetic studies in the fruit fly *Drosophila melanogaster* have led to the identification of highly conserved pathways that regulate both sleep and feeding [48,49,70,71]. Many genes and signaling molecules required for sleep and metabolic regulation are conserved across phyla, including those that regulate the circadian clock, energy stores, and neuropeptide regulation [10,47,72]. For example, leucokinin *(Lk)* and its receptor (*Lkr)* are essential for both postprandial sleep and starvation-induced sleep suppression, suggestion the *Lk* signaling mediates both sleep-feeding interactions [11,47]. Here, we found that targeted knockdown of *Lkr* increases sleep in blood-fed mosquitoes, suggested a wake-promoting role that may be similar to what has been observed for starvation-induced sleep suppression in flies [47]. Additional analysis of the effects of *Lkr* on sleep across different feeding conditions, as well as the generation of stable loss-of-function mutants, will support further dissection of its role in feeding-dependent sleep regulation. In addition, the use of dsRNA to identify regulators of sleep in *Ae. aegypti* provides proof-of-principle for the application of screening approaches in this model that would facilitate the identification of novel sleep regulators.

Our findings raise the possibility that increased sleep and reduced activity following a meal is evolutionarily adaptive. Our findings raise the possibility that increased sleep and reduced activity following a meal is evolutionarily adaptive. There are multiple possible explanations for how these physiological changes might improve survival or reproduction. For example, sleep is proposed to be a mechanism of predator avoidance [73], increasing the likelihood that the female mosquito survives when neither mating nor feeding is required. Second, sleep provides a mechanism of energy conservation that may increase the efficiency of time to the subsequent egg-laying cycle or the likelihood of survival. In the case of *Ae. aegypti*, we found increased sleep and reduced activity for up to 5 days after blood feeding, long after responsiveness to host cues has been restored. This suggests the existence of an opportunistic phase of host-seeking, whereby a bloodmeal provides sufficient nutrients not only for egg production but also to allows *Ae. aegypti* to act more like an ambush predator that can respond to nearby host cues from a chosen resting site without expending substantial energy or increasing predation risk. Once these resources are exhausted, sleep levels decrease and activity increases, corresponding to a more determined host seeking mode, where *Ae. aegypti* may take on greater risks in long-distance host pursuit. Determining whether this effect is specific to *Ae. aegypti* or generalizable across other mosquito species and blood-feeding insects will help inform its adaptive significance.

Together, these results position *Ae. aegypti* as a powerful new model for dissecting how diet shapes sleep. By linking blood feeding and dietary protein to sustained changes in sleep and activity, this work establishes a foundation for exploring the molecular and neural mechanisms that couple feeding state to sleep regulation in a highly specialized dietary context. The availability of sophisticated behavioral assays, alongside emerging genetic tools in *Ae. aegypti*, opens the door to mechanistic studies that can parse the contributions of specific nutrients, signaling pathways, and neuronal circuits to sleep regulation. Looking forward, this system offers an unparalleled opportunity to address broader questions about the adaptive functions of sleep, the evolutionary diversity of sleep–feeding interactions, and the conserved pathways that integrate metabolism and behavior across species.

## MATERIALS AND METHODS

### Mosquito Rearing and Maintenance

All experiments used *Aedes aegypti* mosquitoes of the Liverpool strain (Lvp), originally obtained from Virginia Tech and subsequently maintained in the Adelman laboratory at Texas A&M University under standardized insectary conditions. The *Cyc^−/−^*mutant line was generated as previously described [32] and maintained as a stable homozygous stock in the same facility. Adult mosquitoes were housed in mixed-sex cages at 26–28°C and 70% ± 10% relative humidity under a 15 h light: 9 h dark photoperiod (15 h: 9 h LD). Eggs were surface-sterilized with 2% sodium hypochlorite for 2 min, rinsed thoroughly with deionized water, and hatched in deionized water. Larvae were reared in distilled water and fed daily with finely ground Tetramin® fish food at a standardized, density-dependent regimen. Pupae were individually transferred to mesh-covered emergence cups to ensure accurate tracking of eclosion timing. For blood feeding, mated adult females were offered defibrinated sheep blood (Colorado Serum Company, Denver, CO, USA) through a Parafilm® membrane feeder maintained at 37°C; engorgement was visually confirmed before returning mosquitoes to cages. After blood feeding, females were provided with oviposition substrate (seed germination paper strips in water-filled ovitraps) and allowed a standardized 72-h oviposition period when required. Unless otherwise specified, all experimental assays used mated females aged 3–5 days post-eclosion (dpe).

### Sleep, Activity, and Arousal Measurements

#### Analysis of Postural Changes

Mosquito postural changes were recorded using a two-component imaging system: the first component was an iPhone 16 Pro (Apple Inc., Cupertino, CA, USA), mounted on an adjustable tripod to permit precise control of imaging height and distance; the second component was a custom-built behavioral chamber designed to house animals under controlled light/dark and temperature conditions. The chamber had a white exterior with a cut-out front panel to allow stable camera positioning and unobstructed interior imaging. A flat, non-reflective platform served as the base for sample placement. Mosquitoes were housed individually in transparent Pyrex glass tubes (100 mm in length, 10 mm in diameter; TriKinetics Inc., Waltham, MA, USA), which were secured vertically during standard recordings. Adjustable light sources were mounted in the upper interior of the chamber to provide uniform illumination under a 15 h: 9 h LD cycle. This design minimized external disturbance and maintained a stable environment for behavioral measurements. For experiments involving vibrational stimulation, the glass tube was positioned horizontally on a 3D-printed support platform, with the camera aligned perpendicularly above the sample. Video recordings were analyzed to quantify the interval time between the onset of hind-leg lowering, marking the transition from quiescence, and full contact of the hind leg with the substrate. This interval was operationally defined as the time required to enter a sleep-like state.

#### Evaluation of Arousal threshold using custom-designed DART system

A modified Drosophila Arousal Tracking (DART) system was used for mosquito videography [25]. Briefly, sugar-fed females 3–5 dpe were individually placed in transparent Pyrex glass 100 mm tubes (outside diameter 10 mm and length 100mm; TriKinetics, Waltham, MA, USA) mounted on a vibration-capable platform. The setup was housed in an environmentally controlled chamber as described above for postural analysis. Mosquito behavior was recorded continuously for 24 h using a high-resolution USB camera (Logitech; 800 × 600 resolution; 5 frames/s). To measure arousal threshold, vibratory stimuli of increasing intensity [0.6–1.0 g (acceleration)] were delivered once per hour over one 15 h: 9 h LD cycle, beginning at Zeitgeber time 0 (ZT0). Stimulation, video tracking, and data analysis were performed using a custom MATLAB pipeline as previous described [25]. For activity quantification, videos were subsampled at 1 frame/s, and pixel differences between consecutive frames were calculated relative to a background reference generated from 20 randomly selected frames. Movement was defined as a positional change exceeding a predefined pixel threshold. Immobility periods lasting ≥10 min were classified as sleep bouts. For each stimulus, the duration of ongoing inactivity (i.e., bout length) was recorded immediately before stimulation. A behavioral response was defined as observable movement within 15 s of vibration onset. The proportion of responding mosquitoes was then calculated for each bout-length bin (e.g., 1 min, 3 min, 5 min).

#### Activity Monitoring System

Sleep and locomotor activity were quantified using custom-built infrared activity monitors (LAM10, Trikinetics, Waltham, MA, USA), which detect movement via infrared beam breaks across 3-board stacks. Prior to loading, females were assigned to sugar-fed, blood-fed, or bovine serum albumin (BSA)-fed groups as described below. In brief, mosquitoes were anesthetized with ice and individually loaded into 10-mm-diameter Pyrex glass tubes containing a solidified sugar diet (10% sucrose with 1% agar). Tubes were placed horizontally in the monitoring racks, and beam-break events of each board stack were recorded continuously 6–8 days under controlled environmental conditions. Activity counts were binned into 1-min intervals and beam breaks across 3-board stacks on the same monitor were integrated prior to sleep analysis. Immobility was defined as 10 consecutive minutes without beam breaks, which is consistent with the data collected in analysis of postural changes. Total sleep duration, mean activity level, bout number and sleep duration, and other sleep architecture parameters were computed per individual using custom Python scripts (Supplementary file 1).

#### Video tracking with EthoVision

Video tracking was performed using EthoVision XT 15 (Noldus Information Technology Inc., VA, USA). Individual mosquitoes were transferred into separate wells of a 6-well plate immediately after feeding as previously described in *Drosophila* [33]. Videos were recorded continuously for 24 h (one LD cycle) at 15 frames/s using a USB webcam (LifeCam Studio 1080p HD Webcam, Microsoft). Video acquisition was performed in VirtualDub (v1.10.4). To permit recording during the dark phase while maintaining uniform illumination, the built-in IR-cut filter of the camera was removed and replaced with an infrared long-pass filter (Edmund Optics Worldwide). After acquisition, videos were imported into EthoVision XT 15 to extract x-y coordinate data for each mosquito across the full 24 h recording period. The instantaneous velocity of each tracked subject was calculated and recorded for every frame. Position and velocity data were exported and further analyzed using a custom Perl script (v5.10.0) and Microsoft Excel macros. To classify behavioral state, a velocity threshold of 0.4 mm/s was used, with velocities above 0.4 mm/s defined as wake/activity and velocities below 0.4 mm/s defined as doze/sleep state.

### Sugar, blood, and BSA feeding assay

To assess the effects of feeding status on sleep and locomotor activity, female mosquitoes were fed either defibrinated sheep blood (Colorado Serum Company) or a bovine serum albumin solution (150 mg/mL BSA in PBS supplemented with 1 mM ATP, included as a phagostimulant) using an artificial membrane feeder, and allowed to oviposit for 72 h after feeding when required. Sugar-fed controls were maintained on 10% sucrose and subjected to the same time course as animals provided with blood or BSA. Females that had oviposited were then transferred immediately to the activity monitoring system in 10-mm-diameter Pyrex glass tubes containing a solidified sugar diet (10% sucrose + 1% agar). For EthoVision XT-based assays, mosquitoes were loaded into the tracking system immediately after blood feeding. In all cases, only fully engorged individuals fed with blood or BSA were used for behavioral test, and both experimental and control mosquitoes were briefly anesthetized on ice before loading.

### Double-stranded RNA synthesis and injection

To knock down *AaLKR* (AAEL006636), we used the dsRNA sequence against *AaLKR* reported by Kwon et al., targeting a 382 nt region of the transcript [75]. The dsRNA template was amplified by PCR from adult female cDNA using Phusion DNA polymerase (NEB) and primers containing the T7 RNA polymerase promoter sequence at the 5′ ends. PCR products were verified by agarose gel electrophoresis and purified using the NucleoSpin Gel and PCR Clean-up Kit (Macherey-Nagel). dsRNA was synthesized in vitro using the MEGAscript T7 Transcription Kit (Thermo Fisher) with 500 ng template DNA in an overnight reaction at 37°C. The transcription reaction was treated with RNase-free DNase (TURBO DNase, Thermo Fisher) for 30 min at 37°C and purified using the MEGAclear Transcription Cleanup Kit (Thermo Fisher). Purified dsRNA was verified by agarose gel electrophoresis, quantified by spectrophotometry (NanoDrop One, Thermo Fisher), and stored in aliquots at −80°C until use. dsRNA against EGFP was generated as a negative control. Primer sequences are listed in Table A. Female mosquitoes 3–5 dpe were fed defibrinated sheep blood (Colorado Serum Company) using an artificial membrane feeder, and only fully engorged individuals were retained with males to ensure mating. Approximately 24 h after the blood meal, females were anesthetized on ice and injected into the thorax with dsRNA (1 μg per individual) using a Nanoject II microinjector (Drummond). After injection, mosquitoes were maintained on 30% sucrose for an additional 48 h and provided with a cup lined with damp paper for egg laying. The following day, mosquitoes were loaded into the behavioral monitoring systems.

**Table A.**
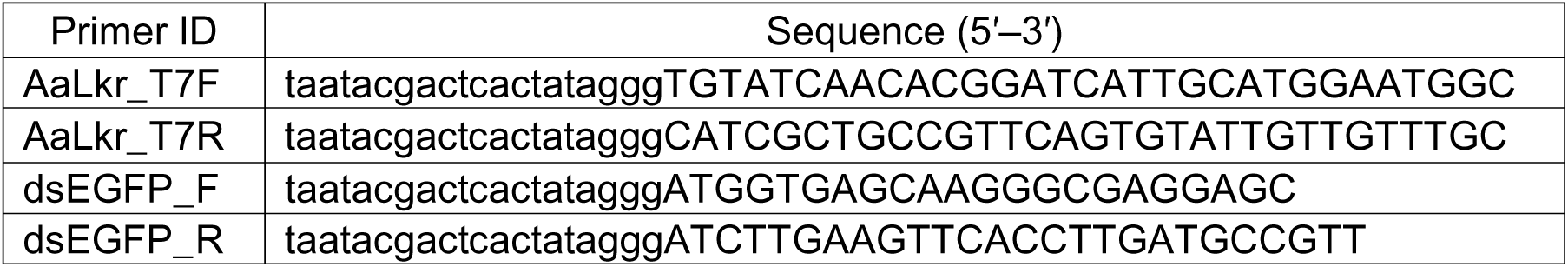
Primers used to amplify dsRNA templates Capital letters indicate gene-specific annealing sequences, and lowercase letters indicate the T7 promoter sequence.

### Statistical analysis

All statistical analyses were performed in GraphPad Prism (version 10.3.0; GraphPad Software, Boston, MA, USA). Data are presented as mean ± SEM unless otherwise indicated. Sample sizes are provided in the figure legends and/or in the Methods. Data distribution and variance were assessed where appropriate before statistical testing. For two-group comparisons, including knockdown experiments comparing mosquitoes injected with dsRNA targeting *AaLkr* or *dsEGFP*, data meeting parametric assumptions were analyzed using unpaired two-tailed t tests, with Welch’s correction applied when variances were unequal. Data that did not meet parametric assumptions were analyzed using Mann–Whitney U tests. For comparisons involving multiple groups, including blood-fed or BSA-fed mosquitoes relative to sugar-fed controls, parametric data were analyzed using one-way ANOVA followed by appropriate multiple-comparison tests, whereas nonparametric data were analyzed using Kruskal–Wallis tests with corrected post hoc comparisons. Paired data shown in paired dot plots were analyzed using paired statistical tests as appropriate. Differences were considered significant at *P* < 0.05. The statistical test used for each analysis is indicated in the corresponding figure legend and supplementary file (Supplementary file 2).

## Data Availability

All data generated or analyzed during this study are included in the manuscript and supporting files; source data file has been provided for all figures. Custom Python scripts for behavioral analysis and statistical analysis spreadsheet have been uploaded as Supplementary file 1 and Supplementary file 2.

---

**Figure 3 figure supplement 1.**
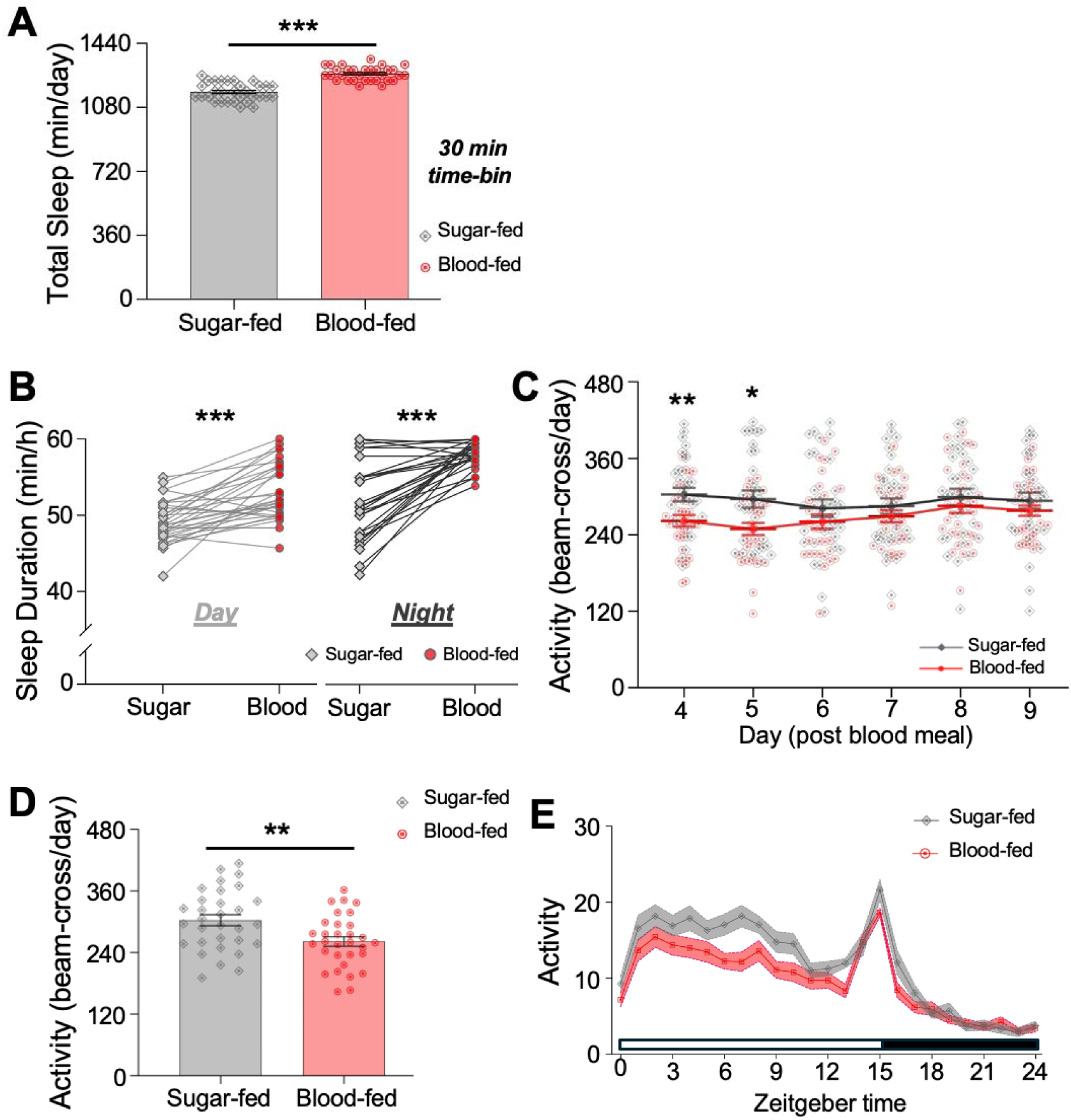
Effects of blood feeding on sleep in *Ae. aegypti* females. (**A**) Total sleep duration in blood-fed Ae. aegypti versus sugar-fed controls in 30-min time-bin of immobility. Sleep of blood-fed mosquitoes (red cycles) was significantly increased compared to sugar-fed controls (gray diamonds, unpaired t-test, *P* < 0.0001). (**B**) Comparisons of the averaged sleep for either sugar-fed or blood-fed individual subjects during daytime and nighttime periods. Two-way ANOVA revealed significant main effects of feeding status (F_1,120_ = 61.60, *P* < 0.0001) and time of day (F_1,120_ = 39.04, *P* < 0.0001), with no significant interaction (F_1,120_ = 2.43, *P* = 0.122). Both sugar-fed and blood-fed mosquitoes slept more during the night than during the day (sugar-fed: paired t-test, *P* = 0.010; blood-fed: paired t-test, *P* < 0.0001). Blood-fed mosquitoes showed significantly higher sleep levels during both daytime and nighttime compared to sugar-fed controls (daytime: unpaired t-test, *P* < 0.0001; nighttime: Mann-Whitney U test, U = 174, *P* < 0.0001). (**C**) Daily activity across Days 4 to 9 post-blood meal. Two-way ANOVA revealed significant effects of time, feeding status, and their interaction (all *P* < 0.0001). Blood-fed mosquitoes (red line) exhibited significantly reduced activity compared to sugar-fed controls (dark gray line) on Days 4 and 5 (Day 4: unpaired t-test, *P* = 0.0045; Day 5: unpaired t-test, *P* = 0.0144), but not on Days 6-9 (Day 6: *P* = 0.337; Day 7: *P* = 0.304; Day 8: *P* = 0.431; Day 9: *P* = 0.431). (**D**) Total daily activity for sugar-fed and blood-fed mosquitoes on Day 4 post blood meal. Blood-fed mosquitoes displayed significantly lower daily activity levels compared to sugar-fed controls (unpaired t-test, *P* = 0.0045). (**E**) Hourly activity profiles across one LD cycle for sugar-fed and blood-fed mosquitoes. Blood-fed mosquitoes showed a distinct reduced activity during both day and night periods. Light-colored shadows indicate the ±SEM error bar, and white and black boxes indicate the daytime and nighttime, respectively. Each dot represents one individual mosquito, dots in gray indicate sugar-fed mosquitoes (N=31) and red dots are for blood-fed animals (N=31). Error bars represent the standard error of the mean (±SEM), and asterisks indicate the level of significance: \**P* < 0.05, \*\**P* < 0.01, \*\*\**P* < 0.001.

**Figure 4 figure supplement 1.**
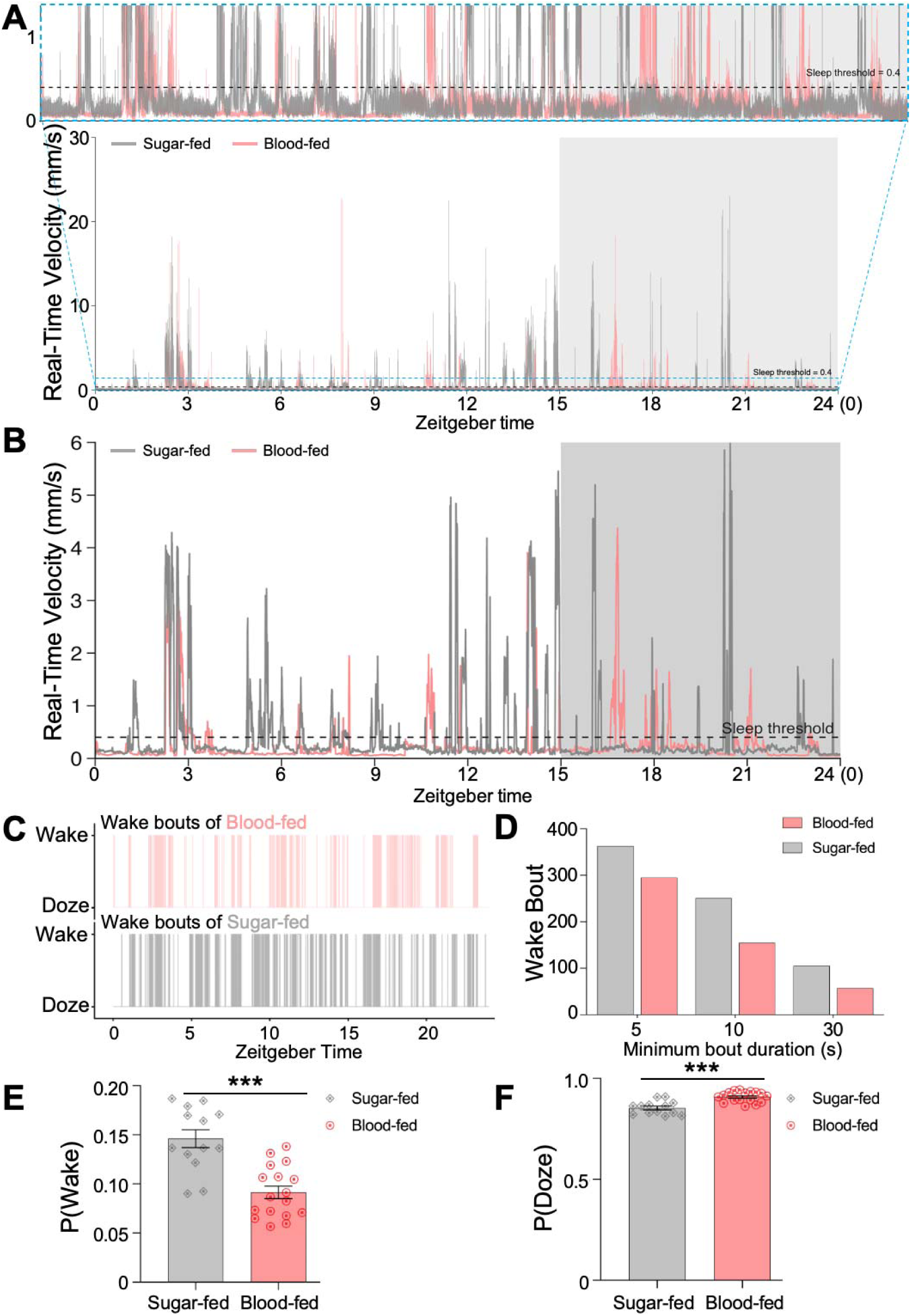
Representative velocity traces and characterization of inactivity/sleep bout duration in sugar-fed and blood-fed female *Aedes aegypti*. (A) Velocity traces (per frame) of sugar-fed and blood-fed female mosquitoes recorded by EthoVision XT. Gray and red traces represent sugar-fed and blood-fed females, respectively. The dashed horizontal line indicates the velocity threshold of 0.4 mm/s used to classify behavioral states: velocity > 0.4 mm/s was defined as wake/activity, and velocity < 0.4 mm/s as doze/sleep. Shaded regions indicate the dark phase. The upper panel shows a zoomed-in view of representative time windows at higher resolution. (**B**) Velocity traces (per second) displayed with rolling smoothing over one second to illustrate locomotor events (wake bouts) in sugar-fed and blood-fed female *Aedes aegypti*. (**C**) Distribution of wake/active bouts in sugar-fed and blood-fed mosquitoes across one light/dark (LD) cycle. (**D**) Number of detected wake/active bouts in sugar-fed and blood-fed mosquitoes using different minimum bout duration thresholds (5 s, 10 s, and 30 s). As expected, the number of detected bouts decreased as the minimum bout duration criterion increased. (**E**) Blood-fed mosquitoes exhibited a significantly lower probability of wakefulness [P(Wake)] compared to sugar-fed controls (unpaired t-test, *P* < 0.0001), consistent with their increased total sleep duration (Figure 4C). Behavioral state occupancy was quantified based on a velocity threshold of 0.4 mm/s: periods with velocity > 0.4 mm/s were classified as wake/activity, whereas periods with velocity < 0.4 mm/s were classified as doze/sleep. (**F**) Blood feeding significantly increased the probability of dozing [P(Doze)] compared to sugar feeding (*P* < 0.0001), inversely mirroring the reduced P(Wake) shown in panel **E**.

